# Microstructural integrity of the major nuclei of the thalamus in Parkinson’s disease

**DOI:** 10.1101/2020.05.21.109660

**Authors:** Nadia Borlase, Daniel Myall, Tracy R. Melzer, Leslie Livingston, Richard Watts, Ross J Keenan, Michael MacAskill, Tim Anderson, John C. Dalrymple-Alford

## Abstract

**Background:** Previous research has shown an association between thalamus and cognition in Parkinson’s disease (PD).

**Objectives:** To investigate the microstructural integrity of the nuclei of the thalamus and relationship with cognition.

**Methods:** Level II Movement Disorder Society Task Force Criteria characterised patients with Parkinson’s disease as cognitively normal (PDN, n=51); with mild cognitive impairment (PD-MCI, n=16) or with dementia (PDD, n=15). Twenty-three healthy control subjects were included for comparison. A *k*-means clustering approach segmented the thalamus into regions representing nine major nuclei. Volume, fractional anisotropy and mean diffusivity of nuclei were compared between cognitive groups and the relationship with cognitive domain z-scores investigated using hierarchical Bayesian regression models.

**Results:** There was an overall progressive increase in mean diffusivity as cognition deteriorated (PDN: 1.4 µm^2^/s (95% uncertainty interval [0.2, 2.7]), PDMCI: 2.4 µm^2^/s [0.8,4.0], PDD: 4.5 µm^2^/s [2.8, 6.3]). The largest increase was in the lateral dorsal nucleus (PDN: 0.3 µm^2^/s [-6.7, 7.2], PDMCI: 5.4 µm^2^/s [-4.7, 16.1], PDD: 14.8 µm^2^/s [5.0, 25.0]). Fractional anisotropy showed minimal change between cognitive groups (PDN: 0.001 [-0.005, 0,007], PDMCI: −0.005 [-0.013, 0.003], PDD: −0.005 [-0.014, 0.003]). Increase in mean diffusivity of the thalamus is associated with a global decline in cognition, the magnitude of the effect was greatest in lateral dorsal nucleus. Fractional anisotropy only showed evidence of a relationship with cognitive domain scores in the lateral dorsal nucleus.

**Conclusions:** The relationship between lateral dorsal nucleus integrity and cognitive changes is likely due to its primary connectivity with frontal and temporal regions.

## Introduction

Parkinson’s disease (PD) is characterised by progressive decline in motor and cognitive function. Up to 83% of patients, conditional on survival, will develop dementia within 20 years of onset (Hely, Reid, Adena, Halliday, & Morris, 2008). The biggest risk of progression to dementia is meeting criteria for PD with mild cognitive impairment (PD-MCI) (Hoogland et al., 2018; Wood et al., 2016). Due to the cognitive profile of PD-MCI being heterogeneous between patients (Dalrymple-Alford et al., 2011; Goldman, Aggarwal, & Schroeder, 2015; Litvan et al., 2012) it is difficult to identify markers of disease progression. Identifying the neuro-correlates of cognitive impairment in PD may aid in understanding the progression to dementia.

The thalamus, centrally located and with reciprocal connectivity from both cortex and brainstem, mediates several communication systems that are associated with the neurodegenerative processes of PD (Haber & McFarland, 2001). Previous DTI studies that have identified the thalamus *a priori* as a region of interest are contradictory. Several have reported no differences between PD and control groups on integrity measures (Chan et al., 2007; Gattellaro et al., 2009; Nicoletti et al., 2006; Rizzo et al., 2008; Schocke et al., 2004), with only two reporting disrupted integrity in the thalamus of PD patients compared to healthy control subjects (Peran et al., 2010; Zhan et al., 2012).

There is some evidence to suggest an overall association between the thalamus and cognition in PD however. In PD patients with normal cognition, atrophy of the thalamus predicted conversion to PD-MCI (Foo et al., 2017) and changes in DTI metrics in the thalamus have been associated with cognitive decline over 12 months in *de novo* patients with sub-clinical cognitive impairment (Zhang et al., 2016). Further, lower thalamic integrity is associated with changes in specific domains of cognition as defined using Movement Disorder Society Taskforce criteria (Litvan et al., 2012) where there was a strong relationship between lower thalamic integrity and decreased attention, executive function and memory (Al Serafy, 2015).

The thalamus was considered as a single region of interest in the above studies, requiring DTI metrics to be averaged across the entire structure. The thalamus is made up of several nuclei, each of which have their own primary regions of connectivity (Behrens, Berg, Jbabdi, Rushworth, & Woolrich, 2007) and each nuclei is targeted at different stages of disease in PD (Braak et al.,2006). Post-mortem studies show neuronal loss restricted to the centromedian/parafascicular complex, for example, while the mediodorsal and anterior principal nuclei are preserved (Henderson, 2000). Examination of the nuclei as separate components may thus provide additional information about the association between the thalamus and cognition in PD.

The unique cyto-architecture of the thalamus supports functionally distinct tasks. Thalamic infarction encompassing anterior and medial regions of the thalamus, for example, is associated with memory deficits, whereas lesions in the posterior and lateral regions have no impact on any aspect of memory (Van der Werf et al., 2003). Functional dissociation is also observable using voxel-based DTI, which allows for identification of localised microstructural changes. Using this method, working memory deficits have been primarily associated with subtle changes to microstructure in only those thalamic nuclei that project to prefrontal and parietal regions and no there was association between working memory and any other thalamic regions in healthy subjects (Piras, Caltagirone, & Spalletta, 2010).

In PD, functional dissociation in the fiber tracts associated with thalamic nuclei has been observed in *de novo* patients. Lower fractional anisotropy (FA), a measure of directionality of white matter tracts, was evident in only those fiber tracts projecting from the anterior nuclei complex (AN), ventral anterior nucleus (VA) and mediodorsal nucleus (MDn) compared to controls. In contrast, white mater pathways connecting posterior nuclei were intact (Planetta et al., 2013).

MR shape analysis of the thalamus supports the hypothesis that specific regions within the thalamus drive the association with cognition. Surface-based volumetric analysis identified atrophic change limited to only the dorsal and anterior regions of the thalamus in PD-MCI patients. Atrophy was an independent predictor of conversion from PD-MCI to PDD and associated with semantic fluency and attention (Chung et al., 2017). To date, the thalamic nuclei have not been isolated in PD and the association between the integrity of nuclei and specific cognitive domain function has not been examined. The present investigation sought to clarify this, defining thalamic regions from DTI data by grouping voxels with the same principal direction of orientation using k-means clustering. This methodology has been successfully applied in healthy control subjects (Behrens et al., 2007; Wiegell, Tuch, Larsson, & Wedeen, 2003) but not yet utilised in a neurodegenerative disease sample. We aimed to use the clustering algorithm to define nine major nuclei of the thalamus by their region of primary connectivity, hypothesising that the thalamus would not show uniform changes but that individual regions would be uniquely associated with cognitive domain changes.

## Methods

### Participant Characteristics

A convenience sample of 103 participants meeting the UK Parkinson’s disease Society’s criteria for idiopathic PD (Hughes, Daniel, Kilford, & Lees, 1992) was recruited from the Movement Disorders Clinic at the New Zealand Brain Research Institute. Initial screening excluded patients with atypical parkinsonian disorder, history of moderate/severe head injury, stroke, early-life learning disability, major psychiatric or medical illness in the previous six months, or without a native level of English. Level of cognitive impairment was defined using Movement Disorder Society Task Force Level II Criteria (Litvan et al., 2012; Woods & Pooley, 2015). PD with dementia (PDD) criteria was performance 2 SD below normative data on two tests in at least two of five MDS cognitive domains (Table 1, Dalrymple-Alford et al., 2011) as well as significant impaired functional activities of daily living. PD with mild cognitive impairment (PD-MCI) was performance 1.5 SD or more below normative data on two tests in at least one cognitive domains. All other patients were classified as Parkinson’s disease without cognitive impairment (PDN). Thirty-two PD participants were drug naïve with respect to anti-parkinsonian medication; in all other PD individuals, all assessments were performed on usual medications. Daily dopaminergic medications were standardized as a levodopa equivalent dose (LED).

**Table 1:** Cognitive and Clinical Status.

|  | HC | PDN | PD-MCI | PDD |
| --- | --- | --- | --- | --- |
| No of participants | 23 | 51 | 16 | 15 |
| Age | 68 (9) | 65 (9) | 69 (9) | 73 (7) |
| Sex M:F | 16:6 | 33:28 | 11:5 | 13:2 |
| Levodopa equiv dose (mg) | - | 230 (310) | 480 (440) | 700 (400) |
| Geriatric depression scale | 0.04 (0.2) | 1.0 (2.0) | 1.5 (2.9) | 3.2 (3.2) |
| Symptom duration (years) | - | 4.8 (3.7) | 9.0 (6.2) | 13.8 (9.1) |
| UPDRS-III | - | 33 (15) | 41 (17) | 65 (19) |
| Hoehn-Yahr | - | 1.9 (0.7) | 2.4 (0.9) | 3.4 (0.8) |
| Montreal Cognitive Assessment | 28 (2) | 27 (2) | 23 (3) | 17 (4) |
| **Cognitive domain scores** |  |  |  |  |
| Attention | 0.5 (0.5) | 0.1 (0.5) | -0.8 (0.5) | -1.8 (0.6) |
| Executive Function | 0.7 (0.6) | 0.3 (0.6) | -0.8 (0.8) | -2.0 (0.5) |
| Learning | 1.0 (0.8) | 0.3 (0.7) | -0.8 (0.5) | -1.6 (0.7) |
| Visuoperception | 0.6 (0.6) | 0.3 (0.5) | -0.7 (0.7) | -1.5 (0.8) |
| Language | 0.1 (0.3) | 0.2 (0.4) | -0.2 (0.7) | -1.5 (0.7) |
| Aggregate Score | 0.7 (0.4) | 0.2 (0.4) | -0.8 (0.3) | -1.8 (0.5) |

Mean (std dev) of clinical and demographic characteristics of subjects. HC = Healthy controls; PDN = PD with normal cognition; PD-MCI = PD with mild cognitive impairment; PDD = PD with Dementia. UPDRS-III = Unified Parkinson's disease Rating Scale-III.

Twenty-nine healthy control subjects, free from a history of major neurological or psychiatric disorders or cognitive impairment were included for comparison. They were selected to match the mean characteristics of the PD group (sex ratio and mean age and years of education years, see Table 1). Neuroradiological screening (R.J.K.) excluded: moderate–severe white matter disease (3 PD, 1 control), extensive white matter abnormality and ventricle abnormality (1 PD), moderate-severe cerebellar infarcts (3 PD, 2 control). Further exclusions were: excessive motion or extreme susceptibility artefacts (5 PD, 2 control); clustering algorithm failure to converge (8 PD, 1 control). The final sample was 82 PD (15 PDD, 16 PD-MCI, 51 PDN), and 23 controls. The research was approved by a Regional Ethics Committee of the New Zealand Ministry of Health and informed consent provided by all participants or their caregiver as appropriate.

### Imaging acquisition and processing

Imaging was performed using a General Electric (GE) 3 Tesla HDxt MRI scanner with an eight-channel head coil. Structural MR images were acquired with a T1-weighted, three dimensional spoiled gradient recalled (SPGR) echo acquisition (TE/TR = 2.8/6.6 ms, TI = 400 ms, flip angle = 15 deg, acquisition matrix = 256 × 256 × 170, slice thickness = 1 mm, FOV = 250 × 250 mm^2^, voxel size 0.98 × 0.98 × 1 mm^3^, scan time = 5 min 6 s). Diffusion images were acquired with a 2D diffusion-weighted, spin echo, echo planar imaging sequence was used to measure microstructural integrity, with diffusion weighting in 28 uniformly distributed directions (b = 1000 s/mm^2^) and 4 acquisitions without diffusion weighting (b = 0 s/mm^2^): TE/TR = 86.4/13000 ms, flip angle = 90 deg, acquisition matrix = 128 × 128 × 48, reconstructed matrix = 256 × 256 × 48, slice thickness = 3 mm, FOV = 240 mm^2^, reconstructed voxel size = 1.88×1.88×3 mm^3^, scan time = 7 min 9 s. Image processing was performed using FSL v4.1.6 (www.fmrib.ox.ac.uk/fsl). Diffusion-weighted images were motion- and eddy current distortion–corrected and rotation of the b matrix. The diffusion tensor was then calculated at each voxel using DTIFIT, producing fractional anisotropy (FA) and mean diffusivity (MD) images, and then brain-extracted using BET. FMRIB’s Integrated Registration and Segmentation Tool (FIRST, v 1.2: Patenaude, Smith, Kennedy, & Jenkinson, 2011), part of FMRIB Software Library (FSL: Jenkinson, Beckmann, Behrens, Woolrich, & Smith, 2012) was used to segment subcortical structures, including the thalamus. Following recommendations of de Jong et al., (2008) a boundary correction of a z-value of 3 was applied to the segmented image, corresponding to 99.98% certainty that voxels belonged to the thalamus. Accuracy was confirmed by visual inspection of all MR images and comparison to anatomical guidelines (Duvernoy, 1991; Nolte, 2007).

### Identification of thalamic nuclei

We aimed to segment the thalamus to identify nine clusters: limbic/association nuclei [Anterior Principal (AP), Lateral Dorsal (LD), Mediodorsal (MDn), Lateral Posterior (LP), Pulvinar (Pu)]; sensorimotor [Ventral Anterior (VA), Ventral Lateral (VL), Ventral Posterior Lateral (VP)] and the non-specific Centromedian/Parafascicular (CM/Pf). A cluster of voxels with similarly orientated diffusion can be assumed to represent a nucleus. Voxels with similar orientation were grouped using a *k-*means clustering algorithm (Wiegell et al., 2003). We hypothesized that the number of clusters as defined by Weigell et al., (2003) (*k*=14) may result in the combination of several smaller nuclei in a neurodegenerative sample. A sub-sample of 5 randomly selected participants tested for the optimal number of clusters prior to completing *k-*means clustering. In this sample only, *k*-means clustering was applied at *k*=[14, 16, 18, and 20] in each subject for each thalamus. Differentiation between lateral nuclei was only visible at *k*=20. Thus, the number of clusters was set at *k*=20 for all participants. Initialisation of the cluster centroids was spatially defined, the center of the first cluster was set as far away from the second cluster as possible, with this practice continuing until all 20 clusters were defined. The distance metric was based on orientation and location of the voxel, ensuring similarly-orientated voxels in close proximity combined to form one cluster. All initial centroids were then regenerated. Convergence was achieved when no voxels changed membership from one cluster to another across iterations.

The 20 clusters were then assigned to one of the nine major nuclei of the thalamus with reference to anatomical guidelines (Duvernoy, 1991; Jones, 2007; Morel, Magnin, & Jeanmonod, 1997). The resulting nuclei (Figure 1) were labelled manually (N.B.). While recognising that each aggregation of clusters is only an approximation of individual thalamic nuclei, they are henceforth labelled by their atlas name. Mean diffusion metrics (FA and MD) were extracted from each of the thalamic nuclei. Each nucleus volume was divided by the individual’s intracranial volume and multiplied by the sample mean intracranial volume to give a normalized volume.

**Figure 1:**
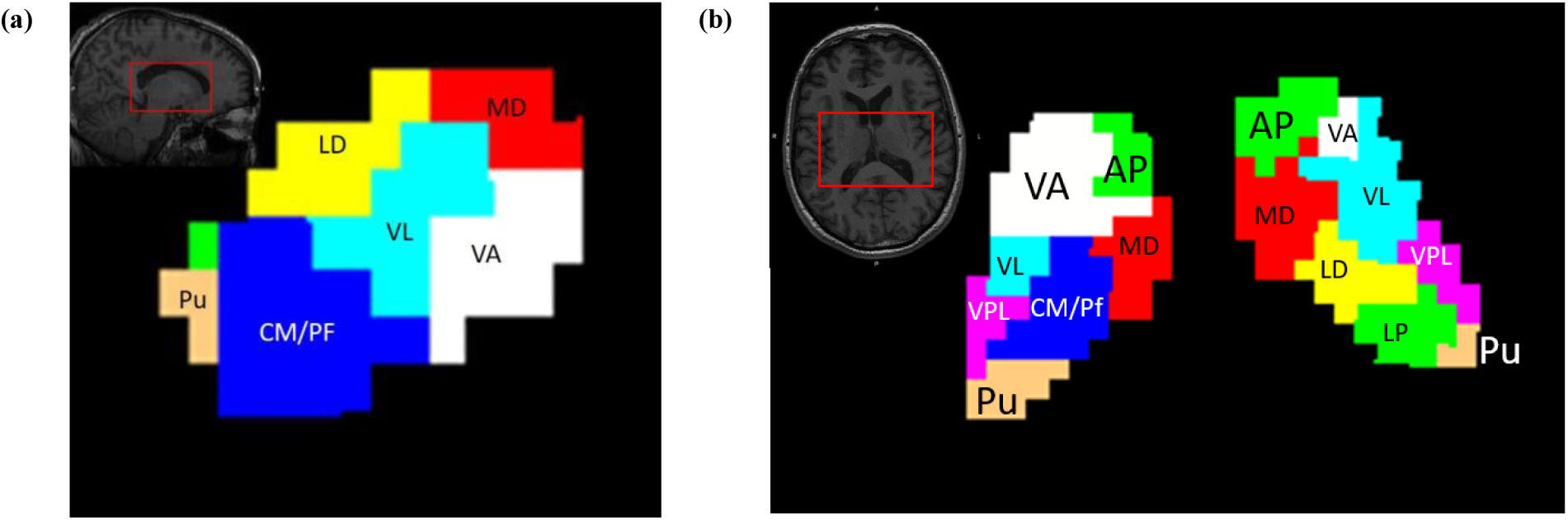
a) sagittal and b) horizontal view of clusters of similarly orientated fibers. Clusters are labelled according to conventional anatomical guidelines overlaid onto T1 weighted image of male PD-MCI patient aged 66 years who was randomly selected for display purposes. Images are displayed in neurological convention where the left side of the image corresponds to the left side of the brain. MNI coordinates: X=70, Y=59, Z=26. NB: The brain is not perfectly straight in this image, thus, in the horizontal image, a more dorsal slice of the thalamus is visible on the right compared to the left.

### Statistical Analysis

Analysis of demographic and cognitive group differences were conducted in R (v 3.3.2, R Core Team, 2019). Bayesian multi-level regression models (Gelman & Pardoe, 2006) investigated the relationship between diffusion measures of thalamic nuclei and group separately for each imaging metric. Models were fit using the “brms” (v1.10.10) package (Burkner, 2017; Carpenter et al., 2017). In each model, four chains with 4000 iterations each generated posterior samples. Models were compared using Pareto-smoothed importance-sampling leave-one-out cross-validation. A higher expected log predictive density leave-one-out (ELPD-LOO) score, by at least five standard errors of the estimated difference, provides strong support for that model over the null model. First, the optimal modelling parameters were determined (see Supplementary Material). Two models were computed to test for a nuclei × group interaction. For each metric, the first model included the dependent variable as a function of the thalamic nuclei only (AP, MD, CM/Pf, LD, VA, VL, VPL, LP, Pu). The second added the group (C, PDN, PD-MCI, PDD) × nucleus interaction factor. For each metric, to determine if the group × nucleus predictor was useful at explaining variance in the data (ie: indicating a group × nucleus interaction) and then the two models were compared. The group × nucleus interaction was then compared to a model which additionally included the covariates of age, sex and Unified Parkinson’s Disease Rating Scale Part III: motor examination (Fahn & Elton, 1987).

## Results

Group demographics and clinical characteristics are summarised in Table 1. The healthy control sample was matched against the mean characteristics of the entire patient sample and, as such, were older than the PDN group, of similar age to PD-MCI and younger than the PDD group. As expected, UPDRS-III motor score, geriatric depression scale (GDS) and levodopa equivalent dose increased from PDN, to PD-MCI, to PDD. By definition, there was a step-wise worsening of cognitive impairment across the PD groups (Z scores PDN > PD-MCI > PDD). The PDN group also generally had poorer cognitive scores than the HC group.

### Between group thalamus differences

#### Volume

The raw data is shown in Figure 2a. There was minimal difference in thalamic nucleus volume between the control group and any PD group (change from control: PDN −1 mm^3^, 95% uncertainty interval [-22, 19]; PD-MCI 4 mm^3^ [-22, 30]; PDD −9 mm^3^ [-34, 18]). In agreement with this, the ELPD-LOO of the model was slightly lower when the group factor was added (delta ELPD-LOO: - 2.1 (SD 1.1)), indicating a slightly worse model fit. Adding age (−16 mm^3^/10 years [-28, −3]) and sex (male −13 mm^3^ [-35, 10]) as covariates slightly improved the model (delta ELPD-LOO: 8.6 (4.9)).

**Figure 2:**
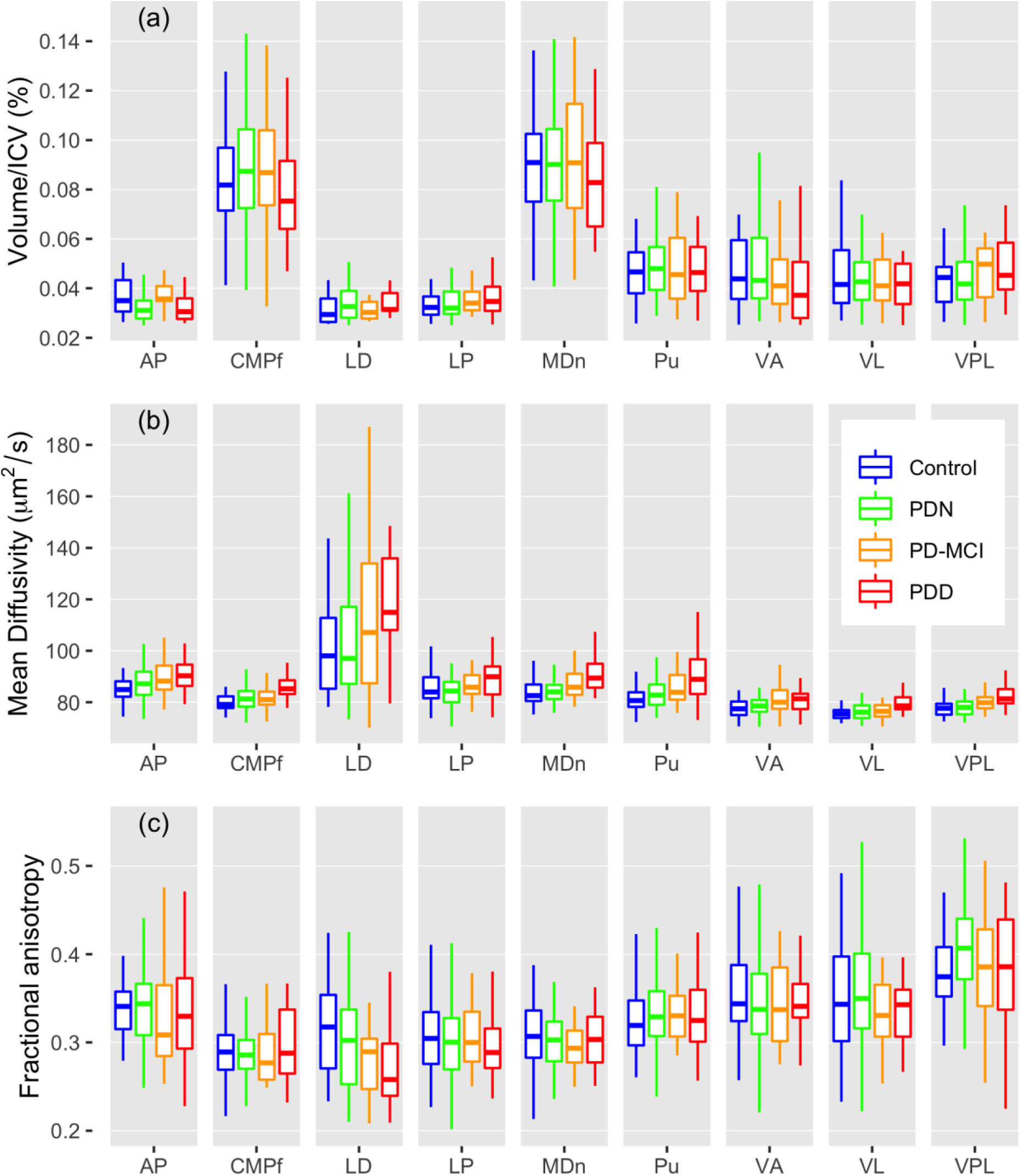
Distribution of (a) Volume, (b) mean diffusivity, and (c) fractional anisotropy by nucleus and cognitive group. The boxes represent the 50th central percentile of the data and the central line the median. The whiskers extend from the box to the value no further than 1.5 times the inter-quartile range.

#### Mean diffusivity

The raw data is shown in Figure 2b. In a model only considering the group effect (collapsing the thalamic nuclei factor), mean diffusivity increased as cognition deteriorated (change from control: PDN: 1.0 µm^2^/s [-0.4, 2.4]; PD-MCI: 2.3 µm^2^/s [0.6,4.1]; PDD: 4.8 µm^2^/s [2.9,6.7]; delta ELPD-LOO=73 (SD 12) compared to a model without group). Adding age and sex as covariates to the model showed both age (1.3 µm^2^/s/10 yrs, [0.8, 2.0]) and being male (−1.3 µm^2^/s [-2.4, −0.1]) were associated with increased mean diffusivity, and the predictive accuracy of the model improved with the addition of these factors (delta ELPD-LOO=70 (SD 10)).

Allowing the effect of group to vary by nuclei resulted in a slight improvement to the model fit (delta ELPD-LOO = 12 (SD 5)). The addition of hemisphere (left, right, by nuclei and group) did not improve the fit (delta ELPD-LOO = −4 (8)). The final model (allowing the effect to vary by nucleus and group, and including covariates) is shown in Figure 3a, where mean diffusivity, relative to controls, had a general pattern of increasing progressively from PDN to PD-MCI to PDD in all nuclei. Across cognitive groups, the sizes of the effects were similar in all nuclei (i.e., a large overlap in uncertainty intervals) except for LD in PDD, where the effect was larger (compared to all other nuclei *n*, probability P_n_ ≥ 95% of a larger effect). Considering mean diffusivity difference in PD-MCI relative to the control group, the size of the MD effect was largest in the limbic/association nuclei: AP P_n_ ≥ 98%; MDn P_n_ ≥ 99%; Pu P_n_ ≥ 99% and the VA P_n_ ≥ 98%, compared to all other nuclei where the effect was P_n_ ≤ 95 %.

**Figure 3:**
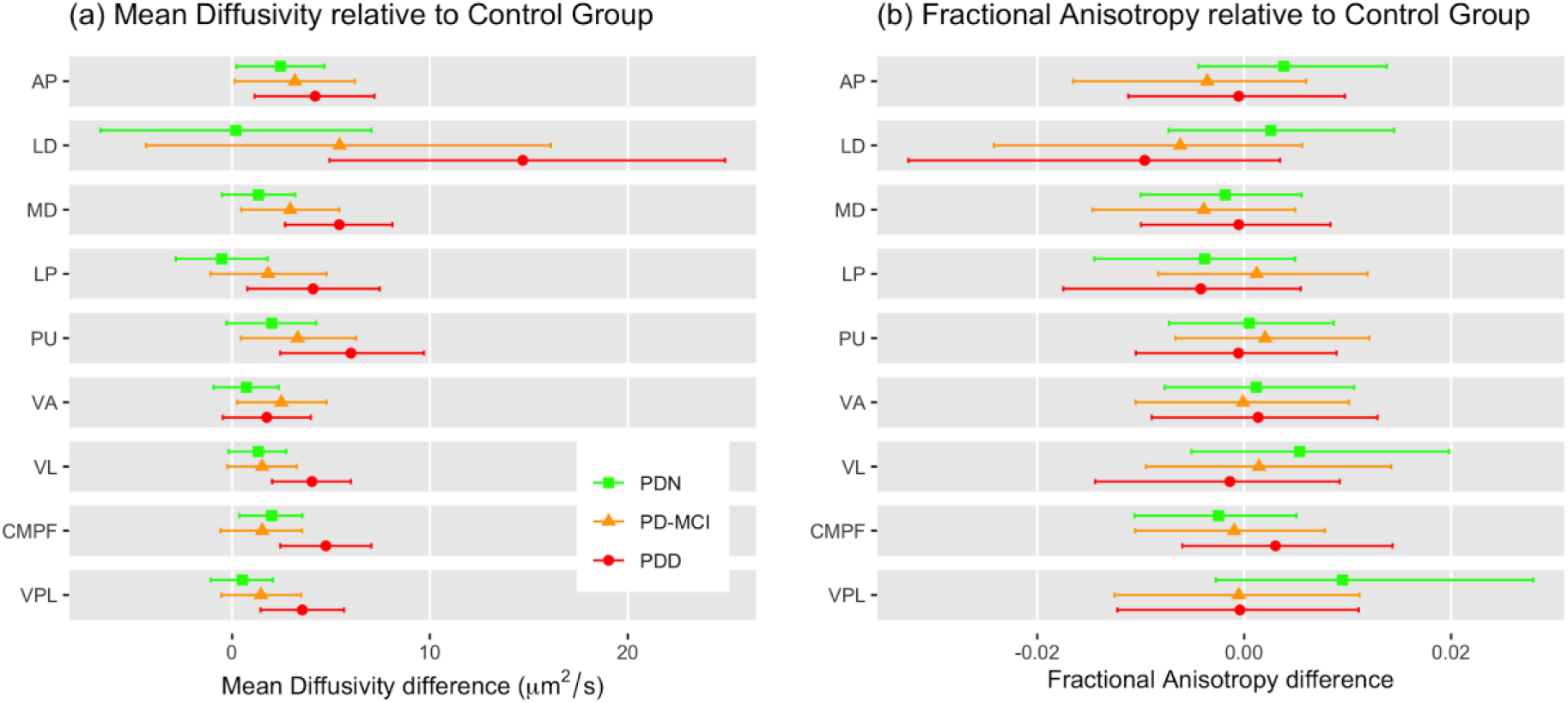
Thalamic nuclei a) mean diffusivity and b) fractional anisotropy differences by group compared to control group. The central points represent the mean of the posterior distribution for that parameter and the error bars the 95% uncertainty intervals **a)** Mean diffusivity: after inclusion of age and sex in the model, there was a general pattern of increased diffusivity from PDN to PD-MCI to PDD, relative to controls, indicating a progressive increase in mean diffusivity as cognition worsened. The greatest difference was evident in the LD nucleus of the PDD group. **b)** Fractional anisotropy: the uncertainty intervals consistently overlap with zero, providing no evidence that there were group differences in fractional anisotropy within thalamic nuclei.

Nuclei differences between groups were not associated with motor impairment, a model including the UPDRS motor score (restricted to patients with UPDRS 3 scores, *n* = 24 patients did not have UPDRS III scores) resulted in a slightly worse model fit (delta ELPD-LOO = −0.8 (0.4)) than the model without UDPRS score.

#### Fractional Anisotropy

The raw data is shown in Figure 2c. There was minimal difference in FA between the control group and any PD group (Figure 3b), (collapsed across nuclei, change from control: PDN −0.001, 95% uncertainty interval [-0.005,0.007]; PD-MCI −0.005 [-0.013,0.003]; PDD −0.005 [-0.014, 0.003]). Correspondingly, the predictive accuracy of the model did not change with the addition of group (delta ELPD-LOO = 1.3(3.0)). Adding age and sex as covariates to the model made a moderate improvement in predictive accuracy of the model (delta ELPD-LOO=20(6.6)). There was no evidence that group effects varied by nuclei, with the addition of this predictor to the model resulting in a slightly worse fit (ELPD-LOO = −3.7±2.7). Although there was a global hemisphere effect (right - 0.0010 [-0.0013, −0.0007]), there was no evidence of variation by group and nucleus (delta ELPD-LOO = −5.4 (5.1)).

### Association with cognition and Parkinson’s disease

#### Mean diffusivity

There was a negative association between mean diffusivity and cognitive domain z-score, as mean diffusivity increased, cognitive score declined (Figure 4a). The degree of association between nuclei and cognition was similar across nuclei (mean −1.8 µm^2^/s per SD of cognition) except the LD where the degree of association was stronger (mean −5.7 µm^2^/s per SD of cognition; In the global domain LD has a stronger association with probability P_n_ ≥ 99% for all other nuclei *n*; attention: P_n_ ≥ 99%; executive function: P_n_ ≥ 99%; language: P_n_ ≥ 94%; learning & memory: P_n_ ≥ 93% visuospatial: P_n_ ≥ 98%).

**Figure 4:**
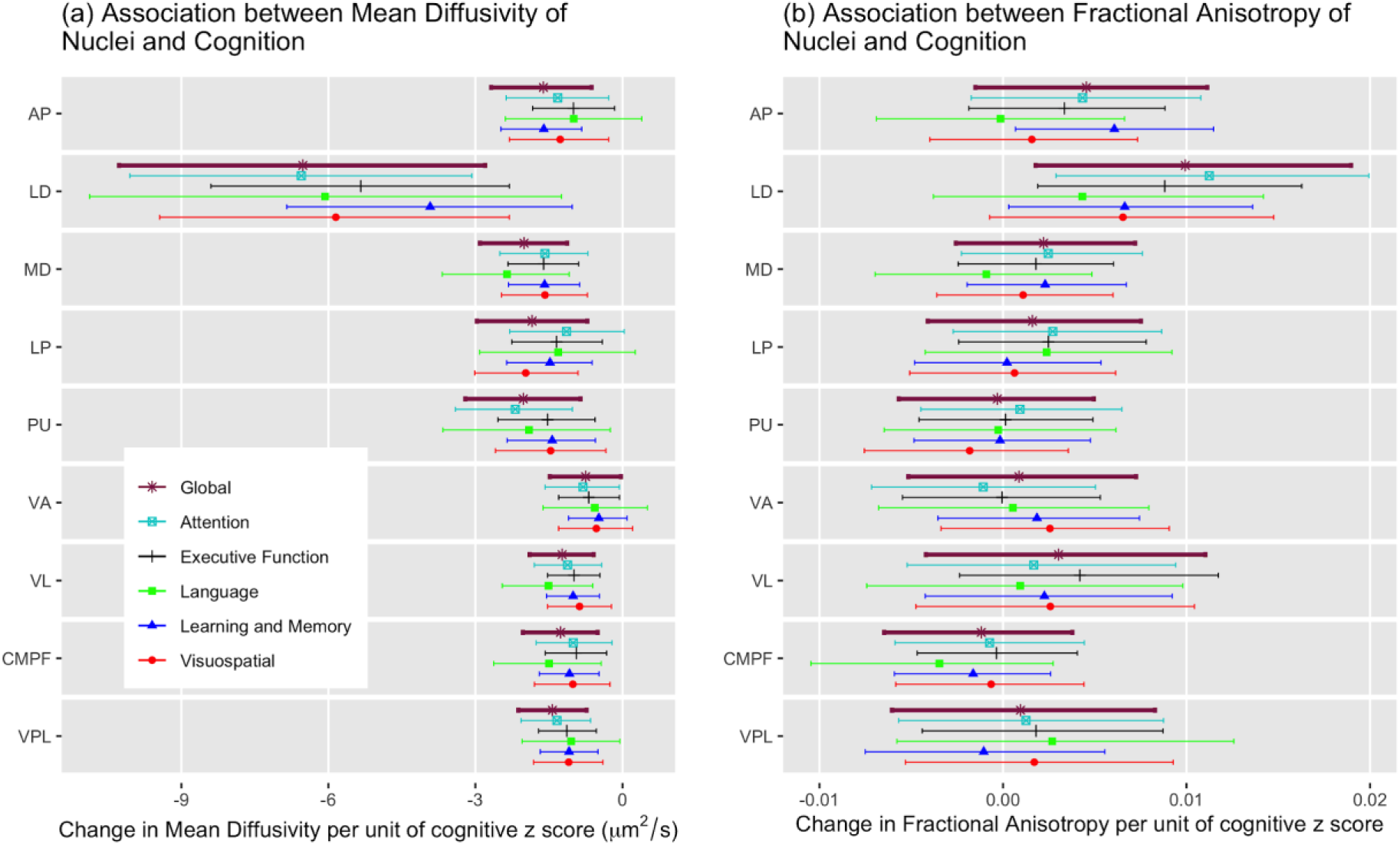
The association between thalamic nucleus integrity and cognition. The central points represent the mean of the posterior distribution for that parameter and the error bars the 95% uncertainty intervals. **a)** In all nuclei there was moderate to strong evidence for an association between mean diffusivity and the cognitive domain score (The probabilities of an association: AP P_d_ ≥ 91% for all domains *d*; LD P_d_ ≥ 99%; MD P_d_ ≥ 99%; LP P_d_ ≥ 95%; Pu P_d_ ≥ 99%; VA P_n_ ≥ 86%; VL P_n_ ≥ 99%; CMPF P_n_ ≥ 99%; VPL P_n_ ≥ 98%). Within each nucleus there was no evidence of differences by cognitive domain. The lateral dorsal (LD) nucleus had the largest association. **b)** There wasn’t strong evidence for a relationship between fractional anisotropy and cognitive domain scores in any nuclei except for LD.

#### Fractional anisotropy

In general, there was minimal association between cognitive z-scores across domains (mean = 0.002 per SD of cognition), with the exception of the LD nucleus, which showed an association between reduced FA and declining cognitive score (mean = 0.008 per SD of cognition) (Figure 4b).

## Discussion

Relative to controls, the thalamus (collapsed across all nuclei) showed a progressive increase in mean diffusivity from PDN to PD-MCI to PDD. Mean diffusivity difference was strongest in the LD, where there was a particularly large difference in PDD compared to controls. There was a moderate increase in all other nuclei except the VA in PDD. In PD-MCI, the limbic and association nuclei (AP, MDn, Pu) and the VA showed the largest difference in mean diffusivity. There was no difference between PDN and controls for any nuclei. In contrast to our hypothesis, DTI measures of thalamic nuclei were similarly associated with each cognitive domain. Mean diffusivity was the most sensitive measure, and associations largest in LD. Fractional anisotropy measures, and the volume of thalamic nuclei, did not show such robust changes by group and there was no relationship between fractional anisotropy or volume of thalamic nuclei with the continuous measures of cognition.

We report a MD increase in the same three nuclei (AP MDn, VA) in PD-MCI identified by Planetta et al., (2013) who reported FA reduction in associated white matter pathways in a *de novo* PD group. Our findings are also supported by Chung et al., (2017) who reported atrophic change in dorsal and limbic regions predicted conversion from PD-MCI to PDD. The limbic/association nuclei and corresponding white matter pathways may therefore be implicated prior to patients meeting criteria for PDD and may play a greater role in cognitive decline in PD than others.

In contrast to our hypothesis, no nuclei were strongly associated with any particular domain. The nuclei of the thalamus therefore appear to be contributing to a global decline in cognition due to involvement in multiple, rather than any single communication pathway. The AP, MDn and VA are integral to maintenance of communication pathways in PD through their role in the limbic; prefrontal and motor loops respectively (Braak & Braak 2000). The stronger role of the LD compared to any other nuclei is likely due to location and connectivity. The LD is located at the point where multiple white matter pathways converge (Perry & Mitchell, 2019). Lesions to the LD disrupt communication with the hippocampus and retrosplenial cortex and result in deficits to attention, spatial memory and visuospatial abilities (Perry & Mitchell, 2019). The association between thalamus and decline in global cognition previously reported (Peran et al., 2010; Zhan et al., 2012) may therefore be driven by association/limbic nuclei, and previous null results (Chan et al., 2007; Gattellaro et al., 2009; Nicoletti et al., 2006; Rizzo et al., 2008; Schocke et al., 2004) due to the effect in these nuclei being washed out when imaging metrics are averaged across the whole thalamus.

Further, DTI is not sensitive to individual white matter pathways within voxels (Glenn et al., 2016), likely to lead to crossing white matter fibers being represented in a single voxel. Regions with crossing fibers may be more susceptible to degeneration than those with linear fiber orientation (Douaud et al., 2011). Thus the location of the LD makes it vulnerable to degeneration.

The raw mean diffusivity measures showed increased variability, particularly within the control group (Figure 2). To account for the deviation from the normal distribution, particularly with respect to outliers, the Bayesian modelling used the Student-t distribution which has increased probability of values falling in the tails and also allowed for the standard deviation within each individual nucleus to be used in the models rather than pooled variance.

A major strength of the present study is the inclusion of a sizeable PD-MCI cohort that was well-characterised according to accepted MDS level II criteria (Litvan et al., 2012), a feature generally missing from previous work (Goldman et al., 2015). In addition, we parcellated the thalamus into specific nuclei; this allowed specific investigation of the relationship between different nuclei-known to exhibit distinct association with the cortex-and different aspects of cognitive impairment. One limitation is the cross-sectional study design and, we plan to follow up the non-demented patients to examine the association between thalamic nuclei microstructural changes as measured with DTI and subsequent conversion to dementia (ie: PDD). In addition, this study employed conventional DTI acquisition and modelling; more advanced diffusion sequences, such as newer implementations of high angular resolution diffusion imaging (HARDI), and associated advanced diffusion models (e.g., constrained spherical deconvolution or diffusion kurtosis imaging) may provide further information about the substructure of the thalamus, its various nuclei, and their relationship with cognitive impairment in PD.

In summary, we have shown that in PD mean diffusivity, relative to controls, increases progressively across cognitive groups from PDN to PD-MCI to PDD in all nuclei with the limbic/association nuclei showing the greatest change in PD-MCI. Importantly the LD nucleus demonstrated the most prominent change in PDD compared to control subject and the strongest association with overall cognition. These findings suggest that mean diffusivity metrics of the thalamus may provide early indicators of cognitive decline and dementia in PD. Longitudinal studies are needed to determine whether that is the case.

