## Supplementary Material for "Microstructural integrity of the major nuclei of the thalamus in Parkinson’s disease"

**Optimal Model Tests**

A Student-t distribution is less sensitive to outliers and thus provides a more robust regression model. To determine if this was an improvement over a Gausssian distribution, models using these two distributions were compared for each metric (mean diffusivity, fractional anisotropy and volume). Each nuclei could potentially have differing variability. To evaluate whether a single pooled variance or multiple variances by nuclei were required, for each metric, a model was then compared where the variance could differ by nucleus, to a model where a common variance was shared across all nuclei.

<u>Volume</u>

There was only minor evidence for the use of a Student-t distribution with ELPD-LOO = 14 (SD 6) over a Gaussian distribution but strong support for a separate variance per nuclei rather than a pooled variance with an ELPD-LOO = 150 (SD 18).

<u>Mean Diffusivity</u>

There was strong evidence for both the use of a Student-t distribution with ELPD-LOO = 850 (SD 90) over a Gaussian distribution, and for a separate variance per nuclei rather than a pooled variance with a further improvement of ELPD-LOO = 290 (SD 30).

<u>Fractional Anisotropy</u>

There was evidence for both the use of a Student-t distribution with ELPD-LOO = 64 (SD 15) over a Gaussian distribution, and for a separate variance per nuclei rather than a pooled variance with a further improvement of ELPD-LOO = 83 (SD 14).
